# The coefficient of determination *R*^2^ and intra-class correlation coefficient from generalized linear mixed-effects models revisited and expanded

**DOI:** 10.1101/095851

**Authors:** Shinichi Nakagawa, Paul C. D. Johnson, Holger Schielzeth

## Abstract

The coefficient of determination *R*^2^ quantifies the proportion of variance explained by a statistical model and is an important summary statistic of biological interest. However, estimating *R*^2^ for generalized linear mixed models (GLMMs) remains challenging. We have previously introduced a version of *R*^2^ that we called *R*^2^_GLMM_ for Poisson and binomial GLMMs, but not for other distributional families. Similarly, we earlier discussed how to estimate intra-class correlation coefficients ICC using Poisson and binomial GLMMs. In this article, we expand our methods to all other non-Gaussian distributions, in particular to negative binomial and gamma distributions that are commonly used for modelling biological data. While expanding our approach, we highlight two useful concepts for biologists, Jensen’s inequality and the delta method, both of which help us in understanding the properties of GLMMs. Jensen’s inequality has important implications for biologically meaningful interpretation of GLMMs, while the delta method allows a general derivation of variance associated with non-Gaussian distributions. We also discuss some special considerations for binomial GLMMs with binary or proportion data. We illustrate the implementation of our extension by worked examples from the field of ecology and evolution in the R environment. However, our method can be used across disciplines and regardless of statistical environments.

## 1. Introduction

One of the main purposes of linear modelling is to understand the sources of variation in biological data. In this context, it is not surprising that the coefficient of determination *R*^2^is a commonly reported statistic because it represents the proportion of variance explained by a linear model. The intra-class correlation coefficient ICC is a related statistic that quantifies the proportion of variance explained by a grouping (random) factor in multilevel/hierarchical data. In the field of ecology and evolution, a type of ICC is often referred to as repeatability *R*, where the grouping factor is often individuals that have been phenotyped repeatedly [1, 2]. We have reviewed methods for estimating *R*^2^and ICC in the past, with a particular focus on non-Gaussian response variables in the context of biological data [2, 3]. These previous articles featured generalized linear mixed-effects models (GLMMs) as the most versatile engine for estimating *R*^2^and ICC (specifically *R*^2^_GLMM_ and ICC_GLMM_). Our descriptions were limited to random-intercept GLMMs, but Johnson [4] has recently extended the methods to random-slope GLMMs, widening the applicability of these statistics (see also, [5, 6]).

However, at least one important issue seems to remain. Currently these two statistics are only described for binomial and Poisson GLMMs. Although these two types of GLMMs are arguably the most popular [7], there are other families of distributions that are commonly used in biology, such as negative binomial and gamma distributions [8, 9]. In this article, we revisit and extend *R*^2^_GLMM_ and ICC_GLMM_ to more distributional families with a particular focus on negative binomial and gamma distributions. In this context, we discuss Jensen’s inequality and two variants of the delta method, which are hardly known among biologists. These concepts are useful not only for generalizing our previous methods, but also for interpreting the results of GLMMs. Furthermore, we refer to some special considerations when obtaining *R*^2^_GLMM_ and ICC_GLMM_ from binomial GLMMs for binary and proportion data, which we did not discuss in the past [2, 3]. We provide worked examples inspired from the field of ecology and evolution, focusing on implementation in the *R* environment [10] and finish by referring to two alternative approaches for obtaining *R*^2^ and ICC from GLMMs along with a cautionary note.

## 2. Definitions of *R*^2^_GLMM_, ICC_GLMM_ and overdispersion

To start with, we present *R*^2^_GLMM_ and ICC_GLMM_ for a simple case of Gaussian error distributions based on a linear mixed-effects model (LMM, hence also referred to as *R*^2^_LMM_ and ICC_LMM_). Imagine a two-level dataset where the first level corresponds to observations and the second level to some grouping/clustering factor (e.g. individuals with repeated measurements) with *k* fixed effect covariates. The model can be written as (referred to as Model 1):

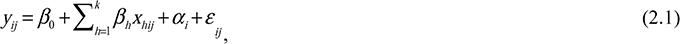

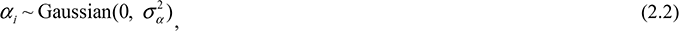

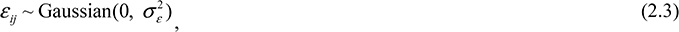

where *y_ij_* is the *j*th observation of the *i*th individual, *x_hij_* is the *j*th value of the *i*th individual for the *h*th of *k* fixed effects predictors, *β*_0_ is the (grand) intercept, *β_h_* is the regression coefficient for the *h*th predictor, *α_i_* is an individual-specific effect, assumed to be normally distributed in the population with the mean and variance of 0 and 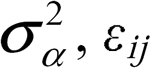 is an observation-specific residual, assumed to be normally distributed in the population with mean and variance of 0 and 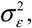 respectively. For this model, we can define two types of *R*^2^ as:

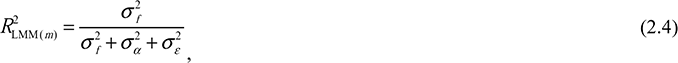

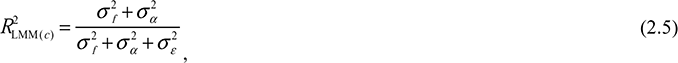

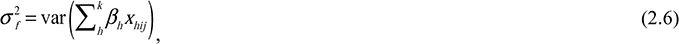

where 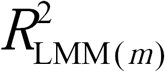 represents the marginal *R*^2^, which is the proportion of the total variance explained by the fixed effects, 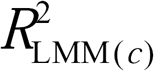 represents the conditional *R*^2^, which is the proportion of the variance explained by both fixed and random effects, and 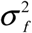 is the variance explained by fixed effects [11]. Since marginal and conditional *R*^2^ differ only in whether the random effect variance is included in the numerator, we avoid redundancy and present equations only for marginal *R*^2^ in the following. Similarly, there are two types of ICC:

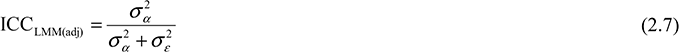

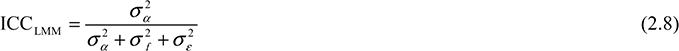

If no fixed effects are fitted (other than the intercept), 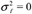 so that ICCLMM(adj) equals ICC_LMM_. In such a case, the ICC should not be called ‘adjusted’ (*sensu* [2]). For an ICC value to be adjusted for a source of variance, that variance must be more than 0 and omitted from the ICC calculation. Since the two versions of ICC differ only in whether the fixed effect variance, calculated as in equation (2.6), is included in the denominator, we avoid redundancy and present equations only for adjusted ICC in the following.

One of the main difficulties in extending *R*^2^ from LMMs to GLMMs is defining the residual variance 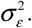 For binomial and Poisson GLMMs with an additive dispersion term, we have previously stated that 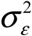 is equivalent to 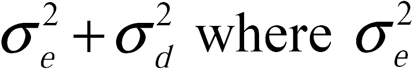 is the variance for the additive overdispersion term, and 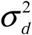 is the distribution-specific variance [2, 3]. Here, overdispersion represents the excess variation relative to what is expected from a certain distribution and can be estimated by fitting an observation-level random effect (OLRE; see, [12, 13]). Alternatively, overdispersion in GLMMs can be implemented using a multiplicative overdispersion term [14]. In such an implementation, we stated that 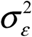 is equivalent to 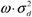 where *ω* is a multiplicative dispersion parameter estimated from the model [2]. However, obtaining 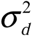 for specific distributions is not always possible, because in many families of GLMMs, 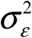 (observation-level variance) cannot be clearly separated into 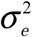 (overdispersion variance) and 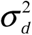 (distribution-specific variance). It turns out that binomial and Poisson distributions are special cases where 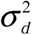 can be usefully calculated, because either all overdispersion is modelled by an OLRE (additive overdispersion) or by a single multiplicative overdispersion parameter (multiplicative overdispersion). This is not the case for other families. However, as we will show below, we can always obtain the GLMM version of 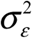 (on the latent scale) directly. We refer to this generalised version of 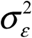 as ‘the observation-level variance’ here rather than the residual variance (but we keep the notation 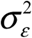). Note that the observation-level variance, 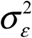, should not be confused with the variance associated with OLRE, which estimates 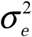 and can be considered to be a part of 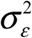.

## 3. Extension of *R*^2^_GLMM_ and ICC_GLMM_

We now define *R*^2^_GLMM_ and ICC_GLMM_ for a quasi-Poisson (may also be referred to as overdispersed-Poisson) GLMM, because the quasi-Poisson distribution is an extension of Poisson distribution [15, 16], and is similar to the negative binomial distribution at least in their common applications [9, 17]. Imagine count data repeatedly measured from a number of individuals with associated data on *k* covariates. We fit a quasi-Poisson (QP) GLMM with the log link function (Model 2):

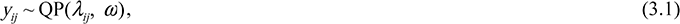

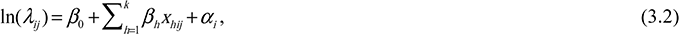

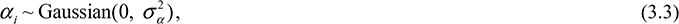

where *y_ij_* is the *j*th observation of the *i*th individual and *y_ij_* follows a quasi-Poisson distribution with two parameters, *λ_ij_* and ω [15, 16], ln(*λ_ij_*) is the latent value for the *j*th observation of the *i*th individual, *ω* is the overdispersion parameter (when the multiplicative dispersion parameter *ω* is 1, the model becomes a standard Poisson GLMM), *α_i_* is an individual-specific effect, assumed to be normally distributed in the population with the mean and variance of 0 and 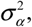, respectively (as in Model 1), and the other symbols are the same as above. Quasi-Poisson distributions have a mean of λ and a variance of *λω* (Table 1). For such a model, we can define 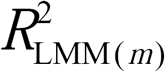 and (adjusted) ICC_GLMM_ as:

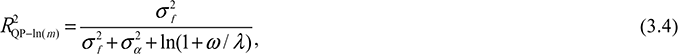

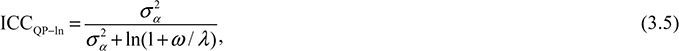

where the subscript of *R*^2^and ICC denote the distributional family, here QP-ln for quasi-Poisson distribution with log link, the term ln(1+*ω* / *λ*) corresponds to the observation-level variance 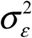 (Table 1, for derivation see Appendix S1), *ω* is the overdispersion parameter, and *λ* is the mean value of *λ_ij_*. We discuss how to obtain *λ* below (Section 5).

**Table 1.**
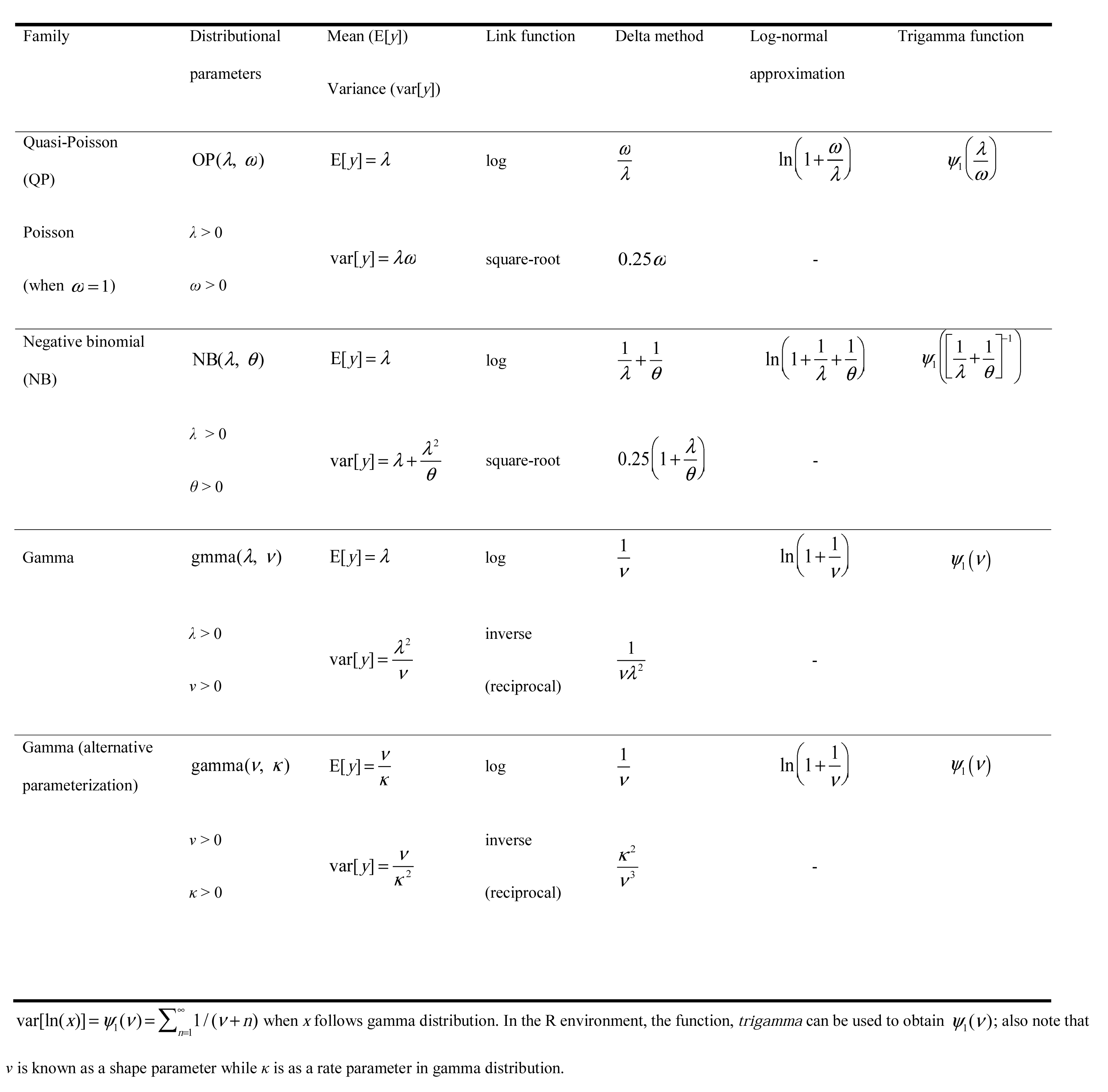
The observation-level variance 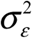 for the three distributional families: quasi-Poisson, negative binomial and gamma with the three different methods for deriving 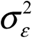: the delta method, log-normal approximation and the trigamma function, *ψ*_1_.

The calculation is very similar for a negative binomial (NB) GLMM with the log link (Model 3):

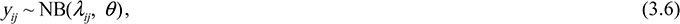

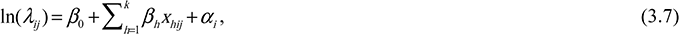

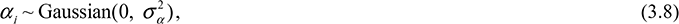

where *y_ij_* is the *j*th observation of the *i*th individual and *y_ij_* follows a negative binomial distribution with two parameters, *λ_ij_* and *θ*, where *θ* is the shape parameter of the negative binomial distribution (given by the software often as the dispersion parameter), and the other symbols are the same as above. The parameter *θ* is sometimes referred to as ‘size’. Negative binomial distributions have a mean of *λ* and a variance of *λ* + *λ*^2^/*θ* (Table 1). *R*^2^_GLMM(*m*)_ and (adjusted) ICC_GLMM_ for this model can be calculated as:

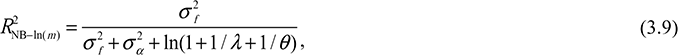

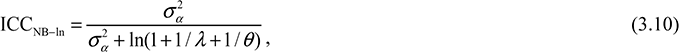

Finally, for a gamma GLMM with the log link (Model 4):

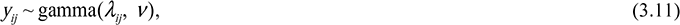

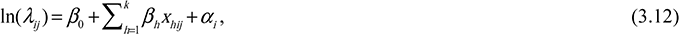

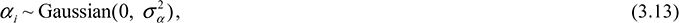

where *y_ij_* is the *j*th observation of the *i*th individual and *y_ij_* follows a gamma distribution with two parameters, *λ_ij_* and *ν*, where *ν* is the shape parameter of the gamma distribution (sometimes statistical programs report 1/*v* instead of *v*; also note that the gamma distribution can be parameterized in alternative ways, Table 1). Gamma distributions have a mean of *λ* and a variance of *λ*^2^/*ν* (Table 1). *R*^2^_GLMM(*m*)_ and (adjusted) ICC_GLMM_ can be calculated as:

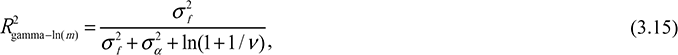

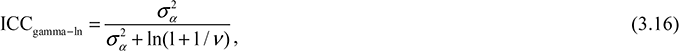

## 4. Obtaining the observation-level variance by the ‘first’ delta method

For overdispersed Poisson, negative binomial and gamma GLMMs with log link, the observation-level variance 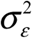 can be obtained via the variance of the log-normal distribution (Appendix S1). This is the approach that has led to the terms presented above. There are two more alternative methods to obtain the same target: the delta method and the trigamma function. The two alternatives have different advantages and we will therefore discuss them in some detail in the following.

The delta method for variance approximation uses a first order Taylor series expansion, which is often employed to approximate the standard error (error variance) for transformations (or functions) of a variable *x* when the (error) variance of *x* itself is known (see [18]; for an accessible reference for biologists, [19]). The delta method for variance approximation can be written as:

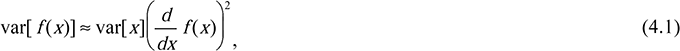

where *x* is a random variable (typically represented by observations), *f* represents a function (e.g. log or square-root), var denotes variance, and *d*/*dx* is a (first) derivative with respect to variable *x*. Taking derivatives of any function can be easily done using the *R* environment (examples can be found in the Appendices). It is the delta method that Foulley and colleagues [20] used to derive the distribution-specific variance 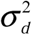 for Poisson GLMMs as 1/*λ* (see also [21]). Given that var[*y*] = *λ* in the case of Poisson distributions and *d* ln(*λ*)/*dx* = 1/*λ*, it follows that var[ln(*y*)] ≈ λ(1/*λ*)^2^ = 1/*λ* (note that for Poisson distributions without overdispersion, 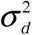 is equal to 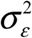 because 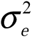 = 0).

One clear advantage of the delta method is its flexibility. We can easily obtain the observation-level variance 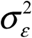 for all kinds of distributions/link functions. For example, by using the delta method, it is straightforward to obtain 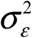 for the Tweedie distribution, which has been used to model non-negative real numbers in ecology (e.g., [22, 23]). For the Tweedie distribution, the variance on the observed scale has the relationship var[*y*] = *φμ^p^* where μ is the mean on the observed scale and *φ* is the dispersion parameter, comparable to *λ* and *ω* in equation (3.1), and *p* is a positive constant called an index parameter. Therefore, when used with the log-link function, 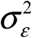 can be approximated by *φμ^(p–2)^* according to equation (4.1). The log-normal approximation ln(1 + *φμ^(p–2)^*) is also possible (see Appendix S1; Table 1).

The use of the trigamma function *ψ*_1_ is limited to distributions with log link, but it is considered to provide the most accurate estimate of the observation level variance 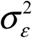 in those cases. This is because the variance of a gamma-distributed variable on the log scale is equal to *ψ*_1_(*ν*) where *ν* is the shape parameter of the gamma distribution [24] and hence 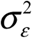 is *ψ*_1_(*ν*). At the level of the statistical parameters (Table 1; on the ‘expected data’ scale; *sensu* [25]; see their Figure 1), Poisson and negative binomial distributions can both be seen as special cases of gamma distributions, and 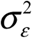 can be obtained using the trigamma function (Table 1). For example, 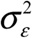 for the Poisson distribution is *ψ*_1_(*λ*) with the speciality that in the case of Poisson distributions 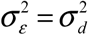. As we show in Appendix S2, ln(1+1/*λ*) (log-normal approximation), 1/*λ* (delta method approximation) and *ψ*_1_(*λ*) (trigamma function) give similar results when *λ* is greater than 2. Our recommendation is to use the trigamma function for obtaining 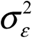 whenever this is possible.

**Figure 1.**
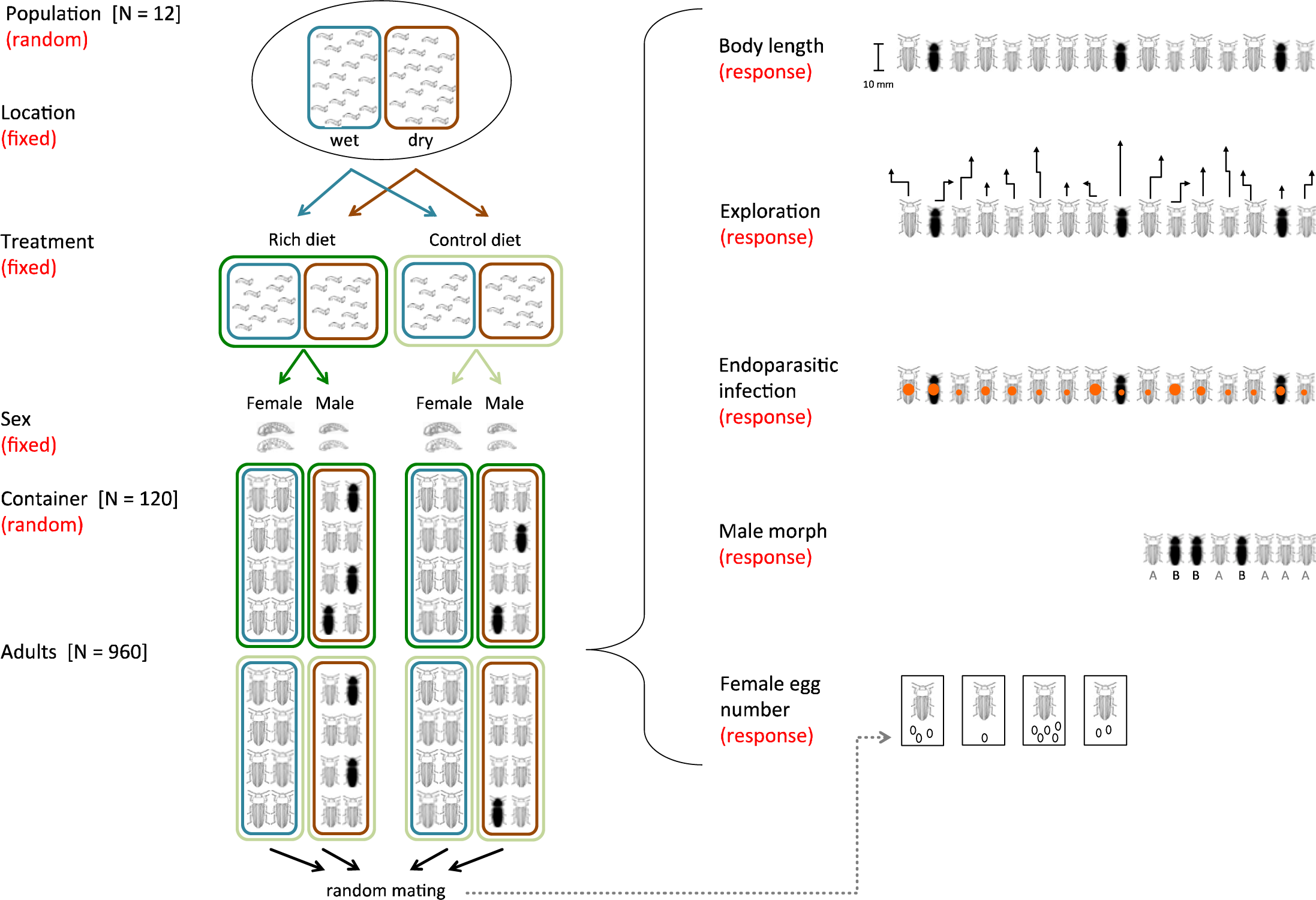
A schematic of how hypothetical datasets are obtained (see the main text for details).

The trigamma function has been previously used to obtain observation-level variance in calculations of heritability (which can be seen as a type of ICC although in a strict sense, it is not; see [25]) using negative binomial GLMMs ([24, 26]; cf. [25]). Table 1 summarises observation-level variance 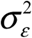 for overdispersed Poisson, negative binomial and gamma distributions for commonly used link functions.

## 5. How to estimate *λ* from data

For some calculations, we require an estimate of the global expected value *λ*. Imagine a Poisson GLMM with log link and additive overdispersion fitted as an observation-level random effect (Model 5):

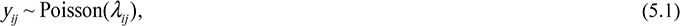

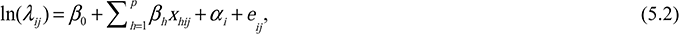

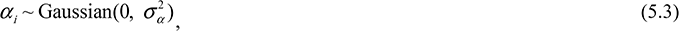

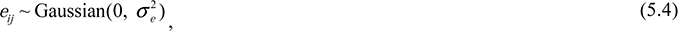

where *y_ij_* is the *j*th observation of the *i*th individual, and follows a Poisson distribution with the parameter **λ*_ij_*, *e_ij_* is an additive overdispersion term for *j*th observation of the *i*th individual, and the other symbols are the same as above. Poisson distributions have a mean of *λ* and a variance of *λ* (cf. Table 1). Using the log-normal approximation *R*^2^_GLMM(*m*)_ and (adjusted) ICC_GLMM_ can be calculated as:

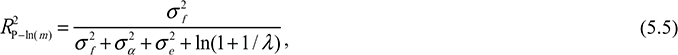

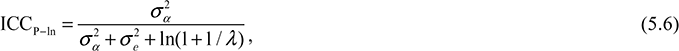

where, as mentioned above, the term ln(1+1/*λ*) is 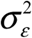 (or 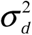) for Poisson distributions with the log link (Table 1).

In our earlier papers, we proposed to use the exponential of the intercept, exp(*β*_0_) (from the intercept-only model) as an estimator of *λ* [2, 3]; note that exp(*β*_0_) from models with any fixed effects will often be different from exp(*β*_0_) from the intercept-only model. We also suggested that it is possible to use the mean of observed values *y_ij_*. Unfortunately, these two recommendations are often inconsistent with each other. This is because, given Model 5 (and all the models in the previous section), the following relationships hold:

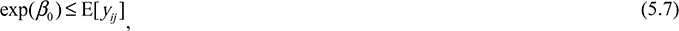

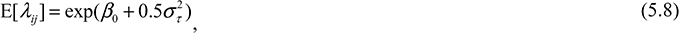

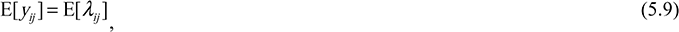

where E represents the expected value (i.e., mean) on the observed scale, *β*_0_ is the mean value on the latent scale (i.e. *β*_0_ from the intercept-only model), 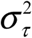 is the total variance on the latent scale (e.g., 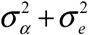 in Models 1 and 5, and 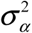 in Models 2-4 [2]; see also [27]). In fact, exp(*β*_0_) gives the median value of *y*_ij_ rather than the mean of *y*_ij_, assuming a Poisson distribution. Thus, the use of exp(*β*_0_) will often overestimate 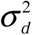, providing smaller estimates of *R*^2^ and ICC, compared to when using averaged *y_ij_* (which is usually a better estimate of E[*y_ij_*]). Quantitative differences between the two approaches may often be negligible, but when *λ* is small, the difference can be substantial so the choice of the method needs to be reported for reproducibility (Appendix S2). Our new recommendation is to obtain *λ* via equation (5.8), which is the Poisson parameter averaged across cluster-level parameters (*λ_i_* for each individual in our example; [[17, 20, 28]]). Thus, obtaining *λ* via equation (5.8) will be more accurate than estimating *λ* by calculating the average of observed values although these two methods will give very similar or identical values when sampling is balanced (i.e., observations are equally distributed across individuals and covariates). This recommendation for obtaining *λ* also applies to negative binomial GLMMs (see Table 1).

## 6. Jensen’s inequality and the ‘second’ delta method

A general form of equation (5.7) is known as Jensen’s inequality, 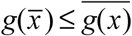 where *g* is a convex function. Hence, the transformation of the mean value is equal to or larger than the mean of transformed values (the opposite is true for a concave function; that is, 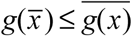; [29]). In fact, whenever the function is not strictly linear, simple application of the inverse link function (or back-transformation) cannot be used to translate the mean on the latent scale into the mean value on the observed scale. This inequality has important implications for the interpretation of results from GLMMs, and also generalized linear models GLMs and linear models with transformed response variables.

Although log-link GLMMs (e.g., Model 5) have an analytical solution, equation (5.8), this is not usually the case. Therefore, converting the latent scale values into observation-scale values requires simulation using the inverse link function. However, the delta method for bias correction can be used as a general approximation to account for Jensen’s inequality when using link functions or transformations. This application of the delta method uses a second order Taylor series expansion [18, 30]. A simple case of the delta method for bias correction can be written as:

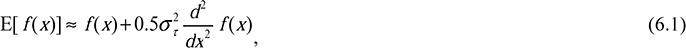

where *d*^2^/*dx*^2^ is a second derivative with respect to the variable *x* and the other symbols are as in equations (4.1) and (5.8). By employing this bias correction delta method (with *d*^2^ exp(*x*)/*dx*^2^ = exp(*x*)), we can approximate equation (5.8) using the same symbols as in equations (5.7)(5.9):

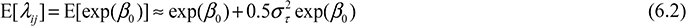

The comparison between equation (5.8) (exact) and equation (6.2) (approximate) is shown in Appendix S3. The approximation is most useful when the exact formula is not available as in the case of a binomial GLMM with logit link (Model 6):

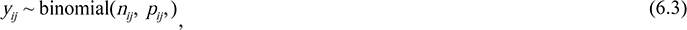

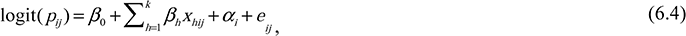

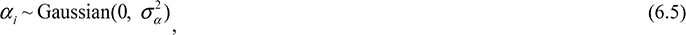

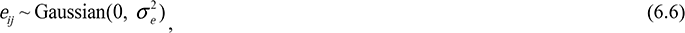

where *y_ij_* is the number of ‘success’ in *n_ij_* trials by the *i*th individual at the *j*th occasion (for binary data, *n_ij_* is always 1), *p_ij_* is the underlying probability of success, and the other symbols are the same as above. Binomial distributions have a mean of *p* and a variance of *np*(1– *p*); (Table 2).

**Table 2.**
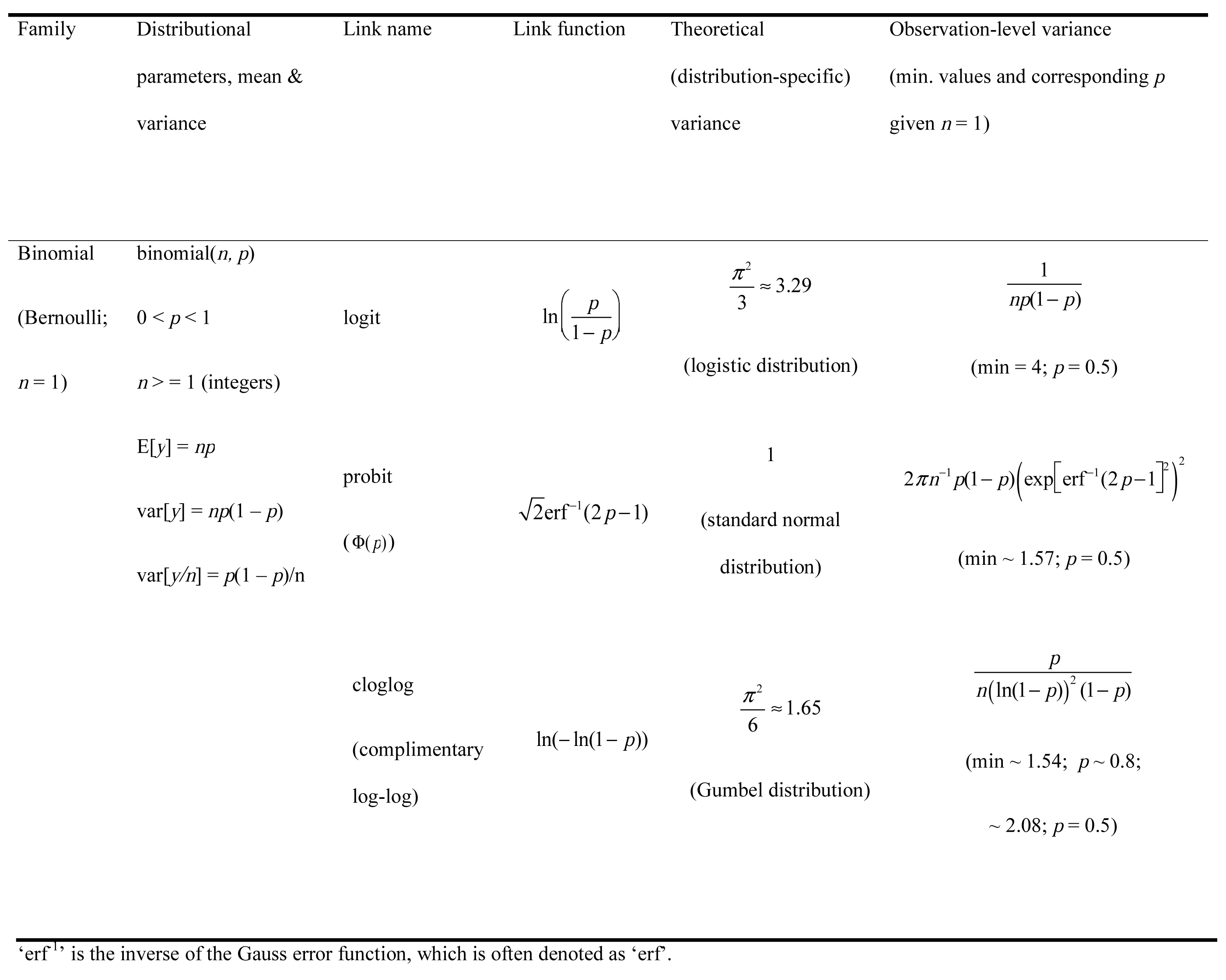
The distribution-specific (theoretical) variance 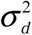 and observation-level variance 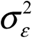 using the delta method for binomial (and Bernoulli) distributions; note that only one of them should be used for obtaining *R*^2^ and ICC.

To obtain corresponding values between the latent scale and data (observation) scale, we need to account for Jensen’s inequality. The logit function used in binomial GLMMs combines of concave and convex sections, which the delta method deals with efficiently. The overall intercept, *β*_0_ on the latent scale could therefore be transformed not with the inverse (anti) logit function (logit^-1^(*x*)/exp(*x*)/(1 + exp(*x*))), but with the bias-corrected delta method approximation. Given that *d*^2^logit^-1^(*x*)/*dx*^2^ = exp(*x*)(1 – exp(*x*)) / (1 + exp(*x*))^3^ in the case of the binomial GLMM with the logit-link function, the approximation can be written as:

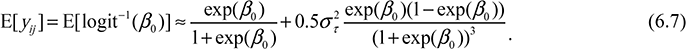

We can replace *β*_0_ with any value obtained from the fixed part of the model (i.e. *β*_0_ + Σ*β_h_x_hij_*. McCulloch and colleagues [31] provide another approximation formula, which, by using our notation, can be written as:

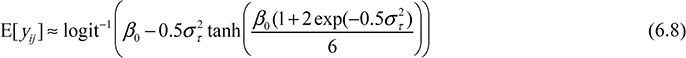

Yet, another approximation proposed by Zeger and colleagues [32] can be written as:

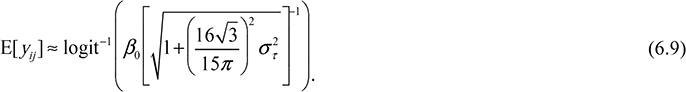

This approximation, equation (6.9), uses the exact solution for the inverse probit function, which can be written for a model like Model 6 but using the probit link: i.e., probit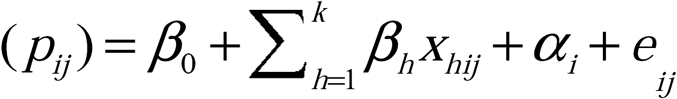in place of equation (6.4)

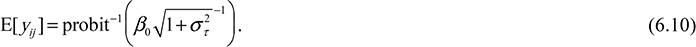

A comparison between equations (6.7), (6.8) and (6.9) is also shown in Appendix S3 (it turns out equation (6.8) gives the best approximation). Simulation will give the most accurate conversions when no exact solutions are available. The use of the delta method for bias correction accounting for Jensen’s inequality is a very general and versatile approach that is applicable for any distribution with any link function (see Appendix S3) and can save computation time. We note that the accuracy of the delta method (both variance approximation and bias correction) depends on the form of the function *f*, the conditions for and limitation of the delta method are described in the article by Oehlert [30].

## 7. Special considerations for binomial GLMMs

The observation-level variance 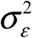 can be thought of as being added to the latent scale on which other variance components are also estimated in a GLMM (equations (3.2), (3.7), (3.12), (5.2) and (6.4) for Models 2-6). Since the proposed *R*^2^_GLMM_ and ICC GLMM are ratios between variance components and their sums, we can show using the delta method that *R*^2^_GLMM_ and ICC GLMM calculated via 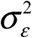 approximate to those of *R*^2^ and ICC on the observation (original) scale (shown in Appendix S4). In some cases, there exist specific formulae for ICC on the observation scale [2]. In the past, we distinguished between ICC on the latent scale and on the observation scale [2]. Such a distinction turns out to be strictly appropriate only for binomial distributions but not for Poisson distributions (and probably also not for other non-Gaussian distributions). This is because the property of what we have called the distribution-specific variance 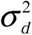 for binomial distributions (e.g. π^2^/3 for binomial error distribution with the logit link function) is quite different from what we have discussed as the observation-level variance 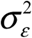 although these two types of variance are related conceptually (i.e., both represents variance due to non-Gaussian distributions with specific link functions). Let us explain this further.

A binomial distribution with a mean of *p* (the proportion of successes) has a variance of *p*(1–*p*)/*n* (the variance for the number of successes is *np*(1–*p*); see Table 2). We find that the observation-level variance is 1/(*np*(1–*p*)) using the delta method on the logit-link function (see Table 2). This observation-level variance 1/(*np*(1–*p*)), or 1/(*p*(1–*p*)) for binary data, is clearly different from the distribution-specific variance π^2^/3. As with the observation-level variance for the log-Poisson model (which is 1/*λ* and changes with *λ*; note that we would have called 1/*λ* the distribution-specific variance; [2, 3]), the observation-level variance of the binomial distribution changes as *p* changes (see Appendix S5), suggesting these two observation-level variances (1/*λ* and 1/(*np*(1–*p*)) are analogous while the distribution-specific variance π^2^/3 is not. Further, the minimum value of 1/(*p*(1–*p*)) is 4, which is larger than π^2^/3 ≈ 3.29, meaning that the use of 1/*p*(1–*p*) in *R*^2^ and ICC for binary data will always produce larger values than those using π^2^/3. Consequently, Browne and colleagues [14] showed that ICC values (or variance partition coefficients, VPCs) estimated using π^2^/3 were higher than corresponding ICC values on the observation (original) scale using logistic-binomial GLMMs (see also [33]). Note that they only considered binary data, i.e., 1/(*np*(1–*p*)) where *n* = 1, because all proportion data can be rearranged as binary responses with a grouping/clustering factor.

Then, what is π^2^/3? Three common link functions in binomial GLMMs (logit, probit and complementary log-log) all have corresponding distributions on the latent scale: the logistic distribution, standard normal distribution and Gumbel distribution, respectively. Each of these distributions has a theoretical variance, namely, π^2^/3, 1 and π^2^/6, respectively, which we previous referred to as distribution-specific variances [2, 3] (Table 2). As far as we are aware, these theoretical variances only exist for binomial distributions. The meaning of 1/(*np*(1–*p*)), which is the variance on the latent scale that approximates to the variance due to binomial distributions on the observation scale is distinct from the meaning of π^2^/3, which is the variance of the latent distribution (i.e., the logistic distribution with the scale parameter being 1). The use of the theoretical variance will almost always provide different values of *R*^2^_GLMM_ and ICC GLMM from those using the observation-level obtained via the delta method (see Appendix S5). This is because the use of π^2^/3 implicitly assumes all data sets have the same observation-level variance regardless of mean (*p*) given the same number of trials (*n*). Therefore, we need distinguishing these theoretical variances from the observation-level variance. *R*^2^ and ICC values using the theoretical distribution-specific variance might be rightly called the latent (link) scale (*sensu* [2]) whereas, as mentioned above, *R*^2^ and ICC values using the observation-level variance estimate the counterparts on the observation (original) scale (cf. [25]).

## 8. Worked examples: revisting the beetles

In the following, we present a worked example by expanding the beetle dataset that was generated for previous work [3]. In brief, the dataset represents a hypothetical species of beetle that has the following life cycle: larvae hatch and grow in the soil until they pupate, and then adult beetles feed and mate on plants. Larvae are sampled from 12 different populations (‘Population’; see Figure 1). Within each population, larvae are collected at two different microhabitats (‘Habitat’): dry and wet areas as determined by soil moisture. Larvae are exposed to two different dietary treatments (‘Treatment’): nutrient rich and control. The species is sexually dimorphic and can be easily sexed at the pupa stage (‘Sex’). Male beetles have two different color morphs: one dark and the other reddish brown (‘Morph’, labeled as A and B in Figure 1). Sexed pupae are housed in standard containers until they mature (‘Container’). Each container holds eight same-sex animals from a single population, but with a mix of individuals from the two habitats (N_[container]_ = 120; N_[animal]_ = 960).

We have data on five phenotypes, two of them sex-limited: (i) the number of eggs laid by each female after random mating which we had generated previously using Poisson distributions (with additive dispersion) and we revisit here for analysis with quasi-Poisson models (i.e. multiplicative dispersion), (ii) the incidence of endo-parasitic infections that we generated as being negative binomial distributed, (iii) body length of adult beetles which we had generated previously using Gaussian distributions and that we revisit here for analysis with gamma distributions, (iv) time to visit five predefined sectors of an arena (employed as a measure of exploratory tendencies) that we generated as being gamma distributed, and (v) the two male morphs, which was again generated with binomial distributions (for the details of parameter settings, see Table 3). We will use this simulated dataset to estimate *R*^2^_GLMM_ and ICC _GLMM_.

**Table 3.** Parameter settings of regression coefficients (*b*) and variance components (σ^2^) for five data sets: 1) fecundity, 2) endoparasite, 3) size, 4) exploration and 5) morph; all parameters are set on the latent scale apart from the size data (see below).

| Response | Intercept ($b$) | Sex ($b$) | Treatment ($b$) | Habitat ($b$) | Population ($\sigma^2$) | Container ($\sigma^2$) | Overdispersion ($\sigma^2$) |
| --- | --- | --- | --- | --- | --- | --- | --- |
| Fecundity: the number of eggs per female | 1.1 | - | 0.5 | 0.1 | 0.4 | 0.05 | 0.1 |
| Parasite: the number of endoparasites per individual | 1.8 | -2 | -0.8 | 0.7 | 0.5 | 0.8 | - |
| Size: the body length of an individual* | 15 | -3 | 0.4 | 0.15 | 1.3 | 0.3 | 1.2 |
| Exploration: the time taken visiting five sectors for an individual | 4 | -1 | 2 | -0.5 | 0.2 | 0.2 | - |
| Morph colour morph of a male | -0.8 | - | 0.8 | 0.5 | 1.2 | 0.2 | - |

\*Data for the six sets of models were simulated on the normal (Gaussian) scale but analysed assuming a gamma error structure with the log link so that estimations of these parameters will be on the log scale; note the overdispersion variance for this data is the residual variance.

All data generation and analyses were conducted in *R* 3.3.1 [10]. We used functions to fit GLMMs from the three *R* packages: 1) the *glmmadmb* function from glmmADMB [34], 2) the *glmmPQL* function from MASS [35], and 3) the *glmer* and *glmer.nb* functions from lme4 [36]. In Table 4, we only report results from *glmmadmb* because this is the only function that can fit models with all relevant distributional families. All scripts and results are provided as an electronic supplement (Appendix S6). In addition, Appendix S6 includes an example of a model using the Tweedie distribution, which was fitted by the *cpglmm* function from the cplm package [23]. Notably, our approach for *R*^2^_GLMM_ is kindly being implemented in the *rsquared* function in the *R* package piecewiseSEM [37]. Another important note is that we often find less congruence in GLMM results from the different packages than those of linear mixed-effects models, LMM. For example, GLMM using the gamma error structure with the log-link function (Size and Exploration models), *glmmadmb* and *glmmPQL* produced very similar results, while *glmer* gave larger *R*^2^ and ICC values than the former two functions (for more details, see Appendix S6; also see [38]). Thus, it is recommended to run GLMMs in more than one package to check robustness of the results although this may not always be possible.

**Table 4.** Mixed-effects model analysis of a simulated dataset estimating variance components and regression slopes for nutrient manipulations on fecundity, endoparasite loads, body length, exploration levels and male morph types; *N*_[population]_=12, *N*_[container]_=120 and *N*_[animal]_=960 (*N*_[male]_= *N*_[female]_ = 480).

| Model name | Fecundity models (log-link) |  | Parasite models (log-link) |  | Size models (log-link) |  | Exploration models (log-link) |  | Morph models (logit-link) |  |
| --- | --- | --- | --- | --- | --- | --- | --- | --- | --- | --- |
|  | Quasi-Poisson mixed models |  | Negative binomial mixed models |  | Gamma mixed models |  | Gamma mixed models |  | Binomial (binary) mixed models |  |
|  | Null Model | Full Model | Null Model | Full Model | Null Model | Full Model | Null Model | Full Model | Null Model | Full Model |
| **Fixed effects** | *b* | *b* | *b* | *b* | *b* | *b* | *b* | *b* | *b* | *b* |
|  | [95% CI] | [95% CI] | [95% CI] | [95% CI] | [95% CI] | [95% CI] | [95% CI] | [95% CI] | [95% CI] | [95% CI] |
| Intercept | 1.630 | 1.261 | 0.766 | 1.752 | 2.682 | 2.737 | 4.752 | 4.056 | -0.108 | -0.740 |
|  | [1.379, 1.882] | [0.989, 1.532] | [0.330, 1.202] | [1.282, 2.223] | [2.616, 2.689] | [2.699, 2.775] | [4.555, 4.949] | [3.842, 4.269] | [-0.718, 0.501] | [-1.450, -0.030] |
| Treatment | - | 0.491 | - | -0.768 | - | 0.033 | - | 2.007 | - | 0.840 |
| (experiment) |  | [0.391, 0.591] |  | [-0.870, -0.667] |  | [0.023, 0.044] |  | [1.965, 2.050] |  | [0.422, 1.258] |
| Habitat (wet) | - | 0.152 | - | 0.700 | - | 0.009 | - | -0.560 | - | 0.414 |
|  |  | [0.055, 0.249] |  | [0.599, 0.801] |  | [-0.001, 0.019] |  | [-0.603, -0.518] |  | [0.002, 0.826] |
| Sex (male) | - | - | - | -2.198 | - | -0.213 | - | -1.105 | - | - |
|  |  |  |  | [-2.511, -1.884] |  | [-0.230, -0.196] |  | [-1.256, -0.955] |  | - |
| **Random effects** | $\sigma^2$ | $\sigma^2$ | $\sigma^2$ | $\sigma^2$ | $\sigma^2$ | $\sigma^2$ | $\sigma^2$ | $\sigma^2$ | $\sigma^2$ | $\sigma^2$ |
| Population | 0.178 | 0.187 | 0.375 | 0.541 | 0.0026 | 0.0039 | 0.071 | 0.104 | 1.002 | 1.111 |
| Container | 0.042 | 0.059 | 1.976 | 0.613 | 0.0140 | 0.0014 | 0.364 | 0.163 | 0.136 | 0.186 |
| Observation-level | 0.477 | 0.349 | 0.873 | 0.397 | 0.0069 | 0.0064 | 1.664 | 0.118 | 4.010 (3.290) | 4.010 (3.290) |
| (Distribution-specific) |  |  |  |  |  |  |  |  |  |  |
| Fixed factors | - | 0.066 | - | 1.479 | - | 0.0116 | - | 1.393 | - | 0.220 |
| $R^2_{\text{GLMM}(m)}$ | - | 9.96% | - | 48.50% | - | 49.54% | - | 78.34% | - | 3.98% (4.57%) |
| $R^2_{\text{GLMM}(c)}$ | - | 46.95% | - | 86.33% | - | 72.52% | - | 93.34% | - | 27.46% |
|  |  |  |  |  |  |  |  |  |  | (31.55%) |
| ICC<sub>[Population]</sub> | 25.33% | 31.30% | 11.53% | 34.44% | 11.38% | 33.17% | 3.40% | 26.94% | 19.48% | 20.95% |
|  |  |  |  |  |  |  |  |  | (22.64%;) | (24.23%) |
| ICC<sub>[Container]</sub> | 5.94% | 9.79% | 60.80% | 39.02% | 59.57% | 12.37% | 17.34% | 42.34% | 2.64% (3.07%;) | 3.50% (4.05%) |
| AIC | 2498.8 | 2412.3 | 4342.6 | 3920.5 | 3379.9 | 3139.5 | 11223.8 | 9004.3 | 605.5 | 589.6 |

95 % CI (confidence intervals) were calculated by the *confint* function in lme4. The observation-level variance was obtained by using the trigamma function. In the Morph models, both the observation-level variance and (theoretical) distribution-specific variance were used; note that ones in brackets use the distribution-specific variance for $R^2$ and ICC. ICC<sub>[Container]</sub> is not a typical ‘repeatability’ but the proportion of variance due to the container effect beyond the population variance.

In all the models, estimated regression coefficients and variance components are very much in agreement with what is expected from our parameter settings (compare Table 3 with Table 4; see also Appendix S6). When comparing the null and full models, which had ‘sex’ as a predictor, the magnitudes of the variance component for the container effect always decrease in the full models. This is because the variance due to sex is confounded with the container variance in the null model. As expected, (unadjusted) ICC values from the null models are usually smaller than adjusted ICC values from the full models because the observation-level variance (analogous to the residual variance) was smaller in the full models, implying that the denominator of, for example, equation (3.5) shrinks. However, the numerator also becomes smaller for ICC values for the container effect from the parasite, size and exploration models so that adjusted ICC values are not necessarily larger than unadjusted ICC values. Accordingly, adjusted ICC_[container]_ is smaller in the parasite and size models but not in the exploration model. The last thing to note is that for the morph models (binomial mixed models), both *R*^2^ and ICC values are larger when using the distribution-specific variance rather than the observation-level variance, as discussed above (Table 4; see also Appendix S4).

## 9. Alternatives and a cautonary note

Here we extend our simple methods for obtaining *R*^2^_GLMM_ and ICC GLMM for Poisson and binomial GLMMs to other types of GLMMs such as negative binomial and gamma. We describe three different ways of obtaining the observational-level variance and how to obtain the key rate parameter *λ* for Poisson and negative binomial distributions. We discuss important considerations which arise for estimating *R*^2^_GLMM_ and ICC GLMM with binomial GLMMs. As we have shown, the merit of our approach is not only its ease of implementation but also that our approach encourages researchers to pay more attention to variance components at different levels. Research papers in the field of ecology and evolution often report only regression coefficients but not variance components of GLMMs [3].

We would like to highlight two recent studies that provide alternatives to our approach. First, Jaeger and colleagues [5] have proposed *R*^2^ for fixed effects in GLMMs, which they referred to as *R*^2^_*β*\*_ (an extension of an *R*^2^ for fixed effects in linear mixed models or *R*^2^_*β*_ by Edwards and colleagues [39]). They show that *R*^2^_*β*\*_ is a general form of our marginal *R*^2^_GLMM_; in theory, *R*^2^_*β*\*_ can be used for any distribution (error structure) with any link function. Jaeger and colleagues highlight that in the framework of *R*^2^_*β*\*_, they can easily obtain semi-partial *R*^2^, which quantifies the relative importance of each predictor (fixed effect). As they demonstrate by simulation, their method potentially gives a very reliable tool for model selection. One current issue for this approach is that implementation does not seem as simple as our approach (see also [40]). We note that our *R*^2^_GLMM_ framework could also provide semi-partial *R*^2^ via commonality analysis (see [41]), since unique variance for each predictor in commonality analysis corresponds to semi-partial *R*^2;^; [42].

Second, de Villemereuil and colleagues [25] have provided a framework with which one can estimate exact heritability using GLMMs at different scales (e.g. data and latent scales). Their method can be extended to obtain exact ICC values on the data (observation) scale, which is analogous to, but not the same as, our ICC GLMM using the observation-level variance, 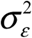 described above. Further, this method can, in theory, be extended to estimate *R*^2^_GLMM_ on the data (observation) scale. One potential difficulty is that the method of de Villemereuil and colleagues is exact but that a numerical method is used to solve relevant equations so one will require a software package (e.g., the QGglmm package; [25]). Relevantly, they have shown that heritability on the latent scale does not need 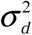 (distribution-specific) but only need 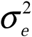 (overdispersion variance), which has interesting consequences in relation to our *R*^2^_GLMM_ and ICC_GLMM_ (we briefly describes this possibility in Appendix S7; see also [40]).

Finally, we finish by repeating what we said at the end of our original *R*^2^ paper [3]. Both *R*^2^ and ICC are indices that are likely to reflect only one or a few aspects of a model fit to the data and should not be used for gauging the quality of a model. We encourage biologists use *R*^2^ and ICC in conjunctions with other indices like information criteria (e.g. AIC, BIC and DIC), and more importantly, with model diagnostics such as checking for model assumptions, heteroscedasticity and sensitivity to outliers.

## Authors’ contribuions

SN conceived ideas, and conducted analysis with discussions with HS. All developed the ideas further, and contributed to writing and editing of the manuscript.

## Acknowledgements

We thank Losia Lagisz for help in making Figure 1. This work has been benefited from discussion with Jarrod Hadfield, Pierre de Villemereuil, Alistair Senior, Joel Pick and Dan Noble. We would also like to thanks an anonymous reviewer, whose comments have improved our manuscript.

